# FrustraMPNN: An ultra-fast deep learning tool for proteome-scale analysis of deep mutational single-residue local energetic frustration in proteins

**DOI:** 10.64898/2026.01.22.701012

**Authors:** Max Beining, Felipe Engelberger, R. Gonzalo Parra, Clara T. Schoeder, César A. Ramírez-Sarmiento, Jens Meiler

## Abstract

Energetic frustration, characterized by conflicting local interactions, is a key determinant of protein dynamics, allostery, protein-protein interactions, enzyme catalysis, and overall protein function. Although the principle of minimal frustration describes an evolutionary bias towards a reduction of energetic conflicts for efficient protein folding, local violations are selectively embedded to encode the complex energy landscapes necessary for protein functions. However, the computational cost of traditional single-residue frustration analysis has prevented the calculation of these energetic conflicts at the proteome scale, creating a significant bottleneck in structural biology. Here, we introduce FrustraMPNN, a message-passing neural network retrained via transfer learning that predicts a complete per-residue frustration-index mutation profile (full mutational scanning per position for a whole protein) orders-of-magnitude faster than existing methods while maintaining high accuracy. This is demonstrated by calculating the single-residue frustration of the *E. coli* proteome, reducing the calculation time from years to 12 hours. We validate FrustraMPNN on a diverse external set of over 3,400 human protein structures, achieving a Spearman correlation of up to 0.80 across all frustration categories, demonstrating robust generalizability. By conducting a thorough dataset ablation study, we find that the model’s performance improves when trained on a broader range of protein sizes, highlighting an inherent limitation of datasets like Megascale. We provide FrustraMPNN as an open-source tool, expecting it will enable exploration of new hypotheses in fields such as personalized structural biology and allow analysis of local energetic frustration patterns involved in protein function at an unprecedented proteomic scale.

## Introduction

Local energetic frustration represents a fundamental principle in protein science, emerging from energy landscape theory and the principle of minimal frustration that governs protein folding [1–3]. In the native state of proteins, most interactions and regions are optimized to maintain structural stability, and these regions are termed minimally frustrated. Mutations in these regions may result in an overall destabilization of the protein structure [4]. Besides that, specific areas with conflicting interactions create local energetic tensions, manifesting as neutral or highly frustrated sites (Figure 1A, top). In contrast to natural proteins, engineered or designed proteins show a different picture as they exhibit an unusually large number of unfrustrated regions due to the primary criteria on minimizing the total energy (Figure 1A, bottom) [1].

**Figure 1.**
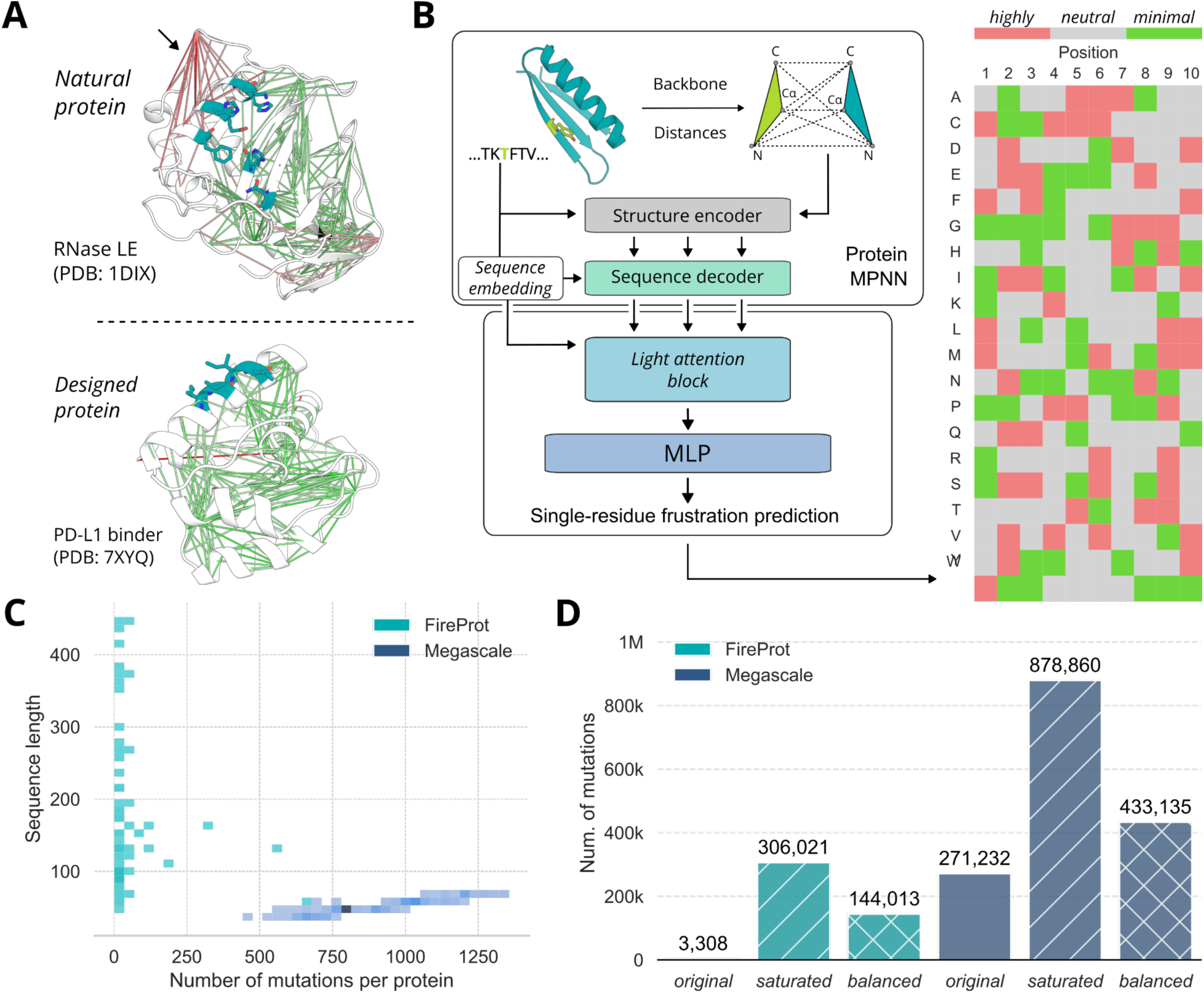
**FrustraMPNN concept and architecture for rapid frustration analysis**. (A) Representative protein structures showing frustration patterns. Top: RNase LE (natural protein) with typical frustration distribution. Bottom: PD-L1 binder (designed protein) showing reduced frustration characteristic of optimized designs. Residues colored by frustration index: green (minimally frustrated), gray (neutral), red (highly frustrated). (B) FrustraMPNN architecture schematic. An input protein structure is encoded as a graph with residue nodes and spatial edges, as in ProteinMPNN. Features pass through message-passing layers into a light attention block and a multilayer perceptron, producing per-residue (single residue) frustration predictions for all positions and all natural amino acids in a single shot. (C) Protein size distribution of the FireProt and Megascale data sets, showing FireProt’s broader coverage and number of mutations per protein and illustrating Megascale’s systematic coverage (deep scan) versus FireProt’s concentrated measurements. (D) Number of total data points/mutations per training set of FireProt and Megascale, split by the original dataset, the saturated set (all possible mutations for all positions), and after balancing for equal frustration category distributions.

Rather than being structural defects, these locally high frustrated regions are often evolutionarily selected features critical for biological function. Frustration patterns have been shown to drive protein dynamics essential for allostery [5], enabling enzyme catalysis by positioning active site residues in energetically unfavorable conformations necessary for chemical reactions [6], mediating protein-protein and protein-ligand binding through conformational flexibility at interfaces [7], and regulating vital biological processes, including viral life cycles and fold-switching behaviors, where frustration governs such conformational changes [1,8,9]. The spatial distribution of local energetic frustration within a protein structure thus provides a physical map of functional importance, making its analysis crucial for understanding protein mechanisms and guiding rational protein engineering and drug discovery efforts [10,11].

It is important to note that energetic frustration alone does not provide information about the function of frustrated regions. It is an indirect metric that provides a way to identify local energetic stresses or disturbances within a protein structure, indicating a certain degree of plasticity or flexibility, but does not reveal the functional outcome. To link local energetic frustration to protein function, it is required to evaluate the direct context of specific chemical or mechanical constraints, either via biophysical experiments to explore the role of frustration in protein dynamics of functionally relevant regions, frustration- and evolution-guided mutagenesis experiments, or comparisons between homologous proteins with divergent frustration patterns.

Notable examples are the frustration analysis, molecular dynamics simulations and nuclear magnetic resonance experiments of apo and active-site-bound thrombin, showing that loop regions surrounding the active site that are highly dynamic in the μs-ms time scale and highly frustrated in apo thrombin are damped upon substrate binding [12,13]. Similarly, the study of variants of enhanced Green Fluorescent Protein (eGFP) generated by random mutagenesis that decreased or increased their folding dependence on the GroEL chaperone system in *E. coli*, showed that mutations that reduced frustration at sites predicted as highly frustrated were less dependent on GroEL for folding [14], thus linking frustration to foldability. Lastly, frustration analysis of the human T4 hormone carrier transthyretin (hTTR) shows that it lost its energetic frustration at the catalytic interface of its enzymatic homolog in *E. coli* (EcTTR), to encode a binding interface that aids in maintaining its structural stability in the bloodstream [15], thus linking the reduction in local frustration to a change in function.

Despite the established importance of frustration in protein biology, its analysis, particularly the comprehensive "single-residue mode" (i.e., the frustration index of all possible mutations in a single residue), remains largely inaccessible to the broader scientific community at scale. The primary barrier is computational: existing physics-based methods, such as those implemented in the Frustratometer web server [16] and FrustratometeR R-package [17], rely on the AWSEM-MD force field [18] to generate and evaluate decoy energies. Deep mutational single-residue frustration analysis is particularly demanding as it requires systematically mutating and energetically evaluating all 20 possible amino acids at each position, not just the native residue or specific mutations. For a 300-residue protein, this calculation requires generating 6,000 structural models – one for every possible substitution – and performing independent frustration calculations for each against a reference ensemble through extensive sampling to achieve statistical significance. This is approximately 20-fold greater than configurational frustration analysis (i.e. comparing the energy of native residue-pair interactions to a statistical baseline of alternative decoy interactions with varying identities, distance and densities of the interacting amino acids), or mutational frustration analysis (i.e. comparing the energy of native residue-pair interactions by generating a set of decoys via randomization of the amino acid identities of the residues in contact). Therefore, single-residue analysis can translate to multiple CPU-days of computation, with larger proteins requiring weeks or even months. The computational cost scales quadratically with protein size because the number of interactions to be evaluated increases, effectively preventing proteome-scale studies in which thousands of proteins require comprehensive single-residue frustration profiling. This means that the analysis of individual amino acid residues – although it is the most meaningful method for identifying functional sites and understanding mutation effects – is limited to small studies of individual proteins.

While critical for function from the perspective of deep mutational scanning, one can place frustrated residues in many places within a protein, but only a small fraction will lead to functional outcomes. The recent revolution in protein science, particularly driven by deep learning models, offers a compelling path forward. Transformative tools such as AlphaFold2 [19] and AlphaFold3 [20] for structure prediction have demonstrated that complex biophysical properties can be learned from data and predicted with remarkable accuracy [21]. In addition, Message-passing neural network (MPNN) architectures have proven particularly effective for protein analysis, with ProteinMPNN achieving state-of-the-art performance in sequence design [22] and ThermoMPNN successfully predicting changes in stability upon mutation [23]. These models leverage the inherent relational graph structure of proteins, where residues form nodes connected by edges representing spatial or chemical relationships, to learn complex patterns from structural data. The success of these approaches in capturing subtle biophysical properties suggests that similar architectures could overcome the computational limitations of single-residue frustration analysis while maintaining the accuracy needed for biological insights.

Here, we present FrustraMPNN, a message-passing neural network developed to overcome the computational limitations of conventional single-residue analysis of local energetic frustration, providing ultra-fast and accurate predictions of complete frustration index profiles for all positions and all mutations of a single protein. Our model achieves orders-of-magnitude speedup compared to existing methods, completing analyses in seconds rather than hours, while maintaining high accuracy across diverse protein structures. We rigorously validate FrustraMPNN on a large external dataset of 3,415 human protein structures predicted by AlphaFold2 available in the AlphaFold Protein Structure Database (AlphaFold DB) [24,25], demonstrating robust generalizability to proteins never seen during training. In addition, this speedup allowed us to analyze the entire *E. coli* proteome (4,362 individual structures) in about 12 h on a single GPU, compared with ∼2 years on a single CPU. We provide FrustraMPNN as a fully open-source tool with comprehensive documentation, thereby democratizing local energetic frustration analysis for researchers across disciplines. Critically, our development and retrospective analysis process revealed that training on a smaller but structurally diverse dataset (FireProt [26]) yielded superior generalizability compared to a larger but more homogeneous dataset (Megascale [27]), highlighting the importance of dataset curation in biological machine learning.

## Results

### FrustraMPNN architecture for high-throughput frustration analysis

FrustraMPNN employs a graph-based neural network (GNN) architecture specifically designed to capture the local structural context that determines local energetic frustration, inspired by the successful protein analysis frameworks ProteinMPNN [22] and ThermoMPNN [23] (Figure 1B). Similar to ProteinMPNN [22], proteins are modeled as graphs, with each residue as a node with edges connecting it to 48 spatially proximate residue neighbors. This graph structure naturally encodes the three-dimensional relationships critical for understanding local energetic conflicts. Then, similar to ThermoMPNN [23], we used the pretrained ProteinMPNN architecture followed by a downstream module to predict single-residue frustration.

Briefly, input backbone coordinates (N, Cα, C, O) and a virtual Cβ are converted into pairwise distances and processed by the structure encoder in ProteinMPNN to capture geometric constraints. The sequence decoder conditions these structural features on the wild-type sequence embedding. Instead of generating autoregressive sequence probabilities, the system extracts the latent embeddings from the decoder. These decoder outputs are concatenated with the original sequence embedding and fed into a Light attention block [28], as used in ThermoMPNN, which assigns weights to different features, enabling the model to focus on the most informative structural characteristics. Finally, a multilayer perceptron (MLP) with two hidden layers maps the learned representations to continuous frustration indices for each residue. Optionally, evolutionary information captured through the ESM-2 language model [29] embeddings are also concatenated to the light attention features and fed into the MLP.

### Dataset characteristics and generalizable prediction on proteome-like diverse external validation

Beyond computational efficiency, a predictive model must demonstrate accuracy and, critically, the ability to generalize over unseen data from distributions different from the training set. We rigorously validated FrustraMPNN using multiple evaluation strategies to ensure robust performance across diverse protein structures and functional classes. Thus, we opted to train FrustraMPNN using two distinct training datasets with contrasting characteristics (Figure 1C and Supplementary Figure S1): (a) the FireProt dataset, based on the FireProtDB database [26] and generated for the development of ThermoMPNN [23] (see Methods), which comprises a total of 3,404 experimentally characterized stability mutations from around 100 natural proteins, representing diverse folds and functions with proteins ranging from 50 to 400 residues; and (b) the Megascale dataset [27] which contains 776,298 systematic mutational scans of natural and de novo designed proteins primarily under 75 residues in length. For the development of ThermoMPNN, this dataset was filtered, resulting in 272,712 mutations across 298 proteins. For FrustraMPNN, single structures from both sets were excluded due to parsing problems with the FrustratometeR [17], resulting in a total mutation count for FireProt of 3,308 and for Megascale of 271,232. While the number of data points obtained from the ThermoMPNN dataset differs, the difference is minimal and should not be critical to our training results.

It is worth noting that both datasets differ not only in the number of data points and protein size, but also in characteristic features, such as the overall proportion of secondary structure and the distribution of mutations, both in terms of mutation types and their locations within the proteins. The original FireProt dataset exhibits a strong alanine bias, meaning that most mutations are to alanine (Supplementary Figure S1 A). At Megascale, however, there is a particular bias toward mutating charged amino acids (lysine K and glutamic acid E) relative to all others (Supplementary Figure S1 B). While most mutations in FireProt are located in the protein core, in Megascale, they are also frequently found on the surface and are therefore more solvent-exposed, as ascertained from the distributions of solvent accessible surface area (SASA) for both datasets (Supplementary Figure S1).

For each dataset, we created three different versions: (a) default (calculating frustration only for mutations present in the FireProt and Megascale datasets); (b) saturated (calculating frustration for all possible mutations at each position for both datasets); and (c) balanced (subsampling to achieve equal representation across minimal, neutral and high frustration categories) (Supplementary Figure S2). As a result, for FireProt the default set contained 3,308 mutations, the saturated set 306,021, and the balanced set 144,013 data points per single-point mutation. For Megascale, the default set contained 271,232 mutations, the saturated set 878,860, and the balanced set 433,135 data points per single-point mutation.

Internal cross-validation using 5-fold splits at the PDB level showed strong performance across all training configurations. Models trained on the balanced FireProt dataset with ESM-2 embeddings [29] and light attention achieved the best internal metrics, with RMSE of 0.716±0.012 and Spearman correlation coefficient of 0.823±0.003 on held-out test proteins using the balanced dataset (Table 1). Notably, while models trained on the larger Megascale dataset showed marginally better performance on their own test splits (RMSE 0.573±0.006), this advantage did not translate to improved generalization.

**Table 1.** Performance comparison of different model configurations on the FireProt and Megascale datasets. Root Mean Square Error (RMSE) and Spearman correlation coefficient (SCC) are reported for models trained using the default, saturated, and balanced training sets to investigate the impact of incorporating light attention and ESM embeddings. Mean and standard deviation are shown for each metric. Bold values indicate the best performance for a given metric and dataset (5-fold cross-validation on different splits) within all datasets and training set categories. (dp - data points per set)

| Dataset | Training set | Add light attention | Add ESM embeddings | RMSE (↓) | SCC (↑) |
| --- | --- | --- | --- | --- | --- |
| FireProt | Default (3,308 dp) | × | × | 0.659 ± 0.034 | 0.699 ± 0.023 |
| | | × | ✓ | 0.639 ± 0.018 | 0.718 ± 0.017 |
| | | ✓ | × | 0.637 ± 0.020 | 0.722 ± 0.015 |
| | | ✓ | ✓ | 0.639 ± 0.023 | 0.725 ± 0.010 |
| | Saturated (306,021 dp) | × | × | 0.660 ± 0.012 | 0.780 ± 0.012 |
| | | × | ✓ | 0.665 ± 0.006 | 0.778 ± 0.012 |
| | | ✓ | × | 0.649 ± 0.014 | 0.790 ± 0.010 |
| | | ✓ | ✓ | 0.654 ± 0.012 | 0.787 ± 0.014 |
| | Balanced (144,013 dp) | × | × | 0.735 ± 0.018 | 0.814 ± 0.005 |
| | | × | ✓ | 0.734 ± 0.021 | 0.814 ± 0.007 |
| | | ✓ | × | 0.721 ± 0.016 | 0.823 ± 0.005 |
|  |  | ✓ | ✓ | **0.716 ± 0.012** | **0.823 ± 0.003** |
| Megascale | Default (271,232 dp) | × | × | 0.654 ± 0.012 | 0.763 ± 0.006 |
| | | × | ✓ | 0.651 ± 0.010 | 0.765 ± 0.004 |
| | | ✓ | × | 0.620 ± 0.014 | 0.793 ± 0.007 |
| | | ✓ | ✓ | 0.625 ± 0.012 | 0.789 ± 0.007 |
| | Saturated (878,860 dp) | × | × | 0.620 ± 0.005 | 0.790 ± 0.003 |
| | | × | ✓ | 0.623 ± 0.005 | 0.788 ± 0.003 |
| | | ✓ | × | **0.573 ± 0.006** | 0.827 ± 0.003 |
| | | ✓ | ✓ | 0.576 ± 0.005 | 0.824 ± 0.002 |
| | Balanced (433,135 dp) | × | × | 0.685 ± 0.006 | 0.836 ± 0.002 |
| | | × | ✓ | 0.688 ± 0.009 | 0.834 ± 0.003 |
| | | ✓ | × | 0.633 ± 0.003 | 0.862 ± 0.002 |
| | | ✓ | ✓ | 0.637 ± 0.004 | **0.859 ± 0.003** |

The actual test of model quality was conducted on an entirely external validation set of 3,415 human protein structures predicted by AlphaFold2 (Supplementary Figure S3). These proteins, with high-confidence predictions (mean pLDDT > 70) and lengths from 50 to 1,000 residues, exhibit a distribution distinct from both training datasets in terms of size, origin, and overall diversity. On this challenging external set, the balanced FireProt-trained model with ESM-2 embeddings and light attention demonstrated superior generalizability, with a weighted-average RMSE of 0.74±0.02 across frustration categories and a Spearman correlation of 0.80±0.01 (Table 2).

**Table 2.** FrustraMPNN performance across training configurations. . The trained models were evaluated on internal test sets (5-fold cross-validation, balanced dataset training) and on the external AlphaFold2 validation set, a random selection of 25 to 1,000 amino acids from the human proteome. Best performance in each category is shown in bold. (RMSE = Root mean squared error, SCC = Spearman correlation coefficient, Acc = Accuracy)

| Model Configuration | Internal Test Set |  |  | External Validation |  |  |
| --- | --- | --- | --- | --- | --- | --- |
|  | RMSE | SCC | Acc. | RMSE | SCC | Acc. |
| Trained on the FireProt dataset |  |  |  |  |  |  |
| Default | 0.659±0.034 | 0.699±0.023 | 0.71 | 0.82±0.04 | 0.68±0.03 | 0.65 |
| Default + ESM | 0.639±0.018 | 0.718±0.017 | 0.73 | 0.79±0.03 | 0.71±0.02 | 0.68 |
| Saturated + LightAtt | 0.649±0.014 | 0.790±0.010 | 0.78 | 0.77±0.03 | 0.75±0.02 | 0.71 |
| Balanced + ESM + LightAtt | 0.716±0.012 | 0.823±0.003 | 0.81 | **0.74±0.02** | **0.80±0.01** | **0.75** |
| Trained on the Megascale dataset |  |  |  |  |  |  |
| Default | 0.654±0.012 | 0.763±0.006 | 0.75 | 0.88±0.05 | 0.65±0.04 | 0.62 |
| Default + ESM | 0.651±0.010 | 0.765±0.004 | 0.76 | 0.86±0.04 | 0.67±0.03 | 0.64 |
| Saturated + LightAtt | **0.573±0.006** | 0.827±0.003 | 0.82 | 0.84±0.04 | 0.69±0.03 | 0.66 |
| Balanced + ESM + LightAtt | 0.637±0.004 | **0.859±0.003** | **0.84** | 0.81±0.03 | 0.72±0.02 | 0.69 |

The FireProt-trained model accurately captured the distribution of single-residue frustration indices, with particularly strong performance in identifying different frustration categories compared to the model trained using the Megascale dataset. Using cutoffs for single-residue minimal frustration indices > 0.55 and < -1 for high frustration indices, based on previous works [4], confusion matrices (Figure 2A) indicate that both models achieve good performance for minimally frustrated regions (FireProt 0.804, Megascale 0.790), but FireProt significantly outperforms Megascale in correctly identifying highly frustrated residues (0.744 vs. 0.654). Both models show a moderate correlation in predicting all frustration indices, with Spearman correlation coefficients of 0.75 for FireProt and 0.73 for Megascale, based on 100 randomly selected structures from the AlphaFold human proteome validation set (Supplementary Figure S4). The overall prediction fraction across all frustration categories does not differ much from that of the calculated categories using FrustratometeR [17], but is highly dependent on the structural context, with the highest Spearman correlation coefficient identified for the protein core and helices as secondary structure elements (Figure 2B,C). The highest prediction accuracy (F1-score ≥ 0.75) is observed for Gly, Ile, Val, Thr, Ser, Leu, Cys, and His, with values between ∼0.78 and ∼0.90. An intermediate performance level (0.75 ≥ F1-score ≥ 0.60) includes Phe, Arg, Asp, Lys, Gly, Asn, Gln, and Ala. The group with the lowest performance (F1-score < 0.60) consists of Pro, Met, Tyr, and Trp, with Trp having the lowest value at ∼0.50 (Figure 2D).

**Figure 2.**
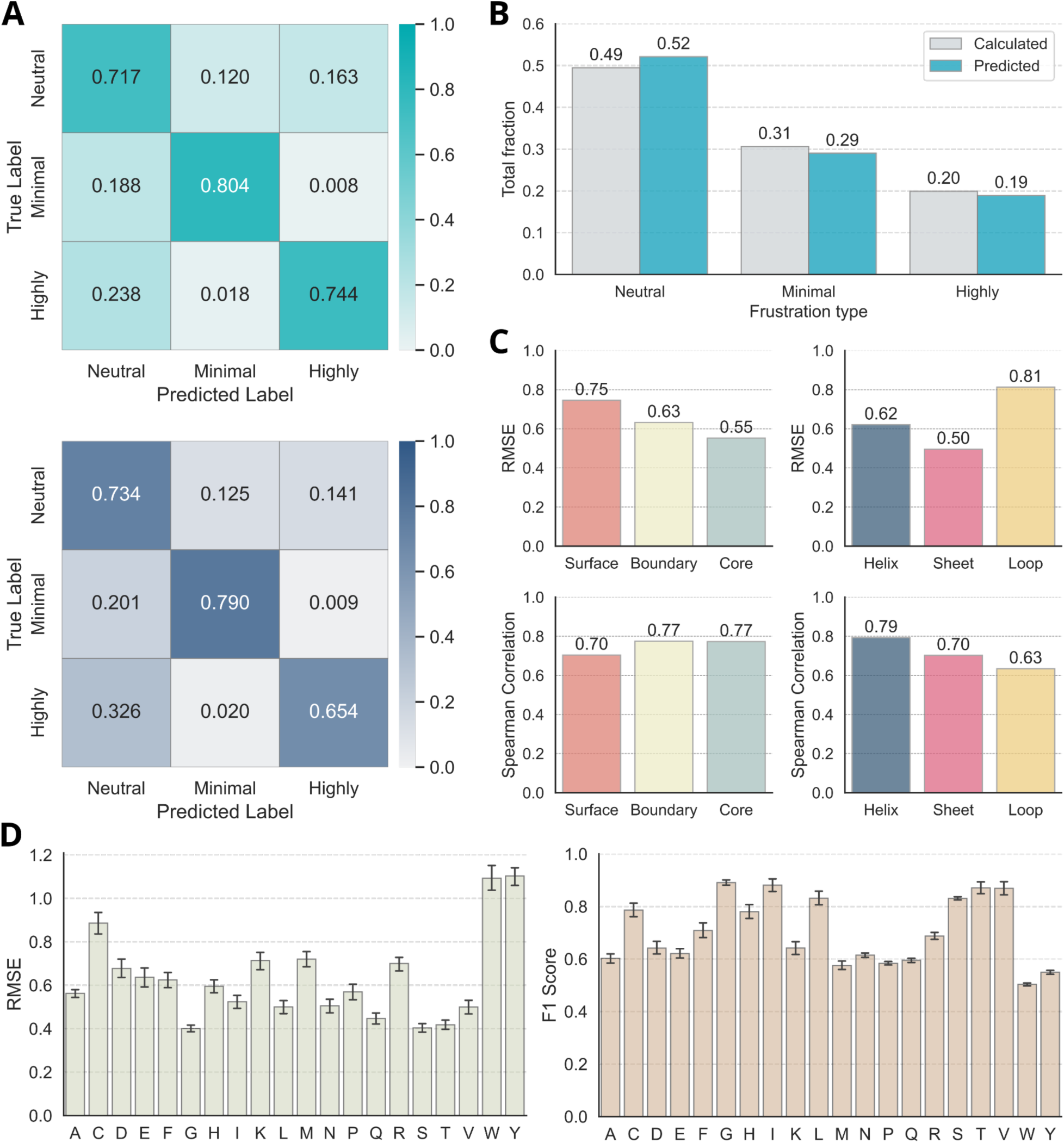
Prediction quality for the validation set for residue location and identity. (A) Confusion matrices comparing classification performance for frustration categories (Neutral, Minimal, Highly) on an external human proteome validation set using models trained on FireProt (top) and Megascale (bottom) datasets. Diagonal entries represent category-specific recall, indicating the FireProt model achieves higher accuracy for highly frustrated residues (0.744) relative to the Megascale model (0.654) (see Tables 1 and 2). (B) Comparison of the global distribution of frustration classes (Neutral, Minimal, Highly) between calculated ground truth (grey) and predictions (teal) for the prediction using the best FireProt model (see Table 2). (C) Predictive accuracy stratified by structural context. Left panels show RMSE and Spearman correlation coefficients across residue solvent accessibility (Surface, Boundary, Core). The right panels show validation metrics across secondary structure elements (Helix, Sheet, and Loop). (D) Amino acid-specific performance (RMSE, F1-score) analysis for single-residue frustration prediction per amino acid type.

### FrustraMPNN matches local protein frustration in enzyme active sites

To assess the predictive capabilities of FrustraMPNN, we compared its output against structure-based calculations from FrustratometeR across three representative enzymes: Class A beta-lactamase (PDB: 4BLM), Serine-carboxyl proteinase (PDB: 1NLU), and Beta-glucosidase (PDB: 1CBG). Visual inspection of the structural projections confirms that FrustraMPNN accurately recapitulates the global frustration landscapes, particularly preserving the high-frustration signals characteristic of catalytic sites (Figure 3). For example, in beta-lactamase, the highly frustrated key residues Lys73, Glu166, and Lys234, together with the neutrally frustrated Ser70, Ser130, and Ala237, were consistently identified by FrustratometeR calculations and FrustraMPNN predictions (Figure 3A), which also agrees with the annotation from previous works [6]. The comparison of the total number of predicted categories revealed a slight shift from 24 (calculated) to 21 highly frustrated sites, from 81 to 76 minimal frustrated, and from 151 to 159 neutrally frustrated residues. Of the initially neutral residues, a small proportion were classified as highly and minimally frustrated. It also indicates that discrepancies are sparse and primarily restricted to transitions between neutral and minimal states, rather than significant misclassifications of highly frustrated functional residues. The incorrectly classified positions are scattered throughout the entire protein. Quantitative analysis via Sankey diagrams showed strong conservation of frustration states for beta-lactamase (Figure 3A, top right). When examining the beta-lactamase in terms of its secondary structure, the most common errors in prediction are found in helices and loops, with a higher proportion of misclassified loop residues located on the surface. Misclassified residues in helices occur equally frequently on the surface than in the core (Figure 3A, bottom and Supplementary Table S1).

**Figure 3.**
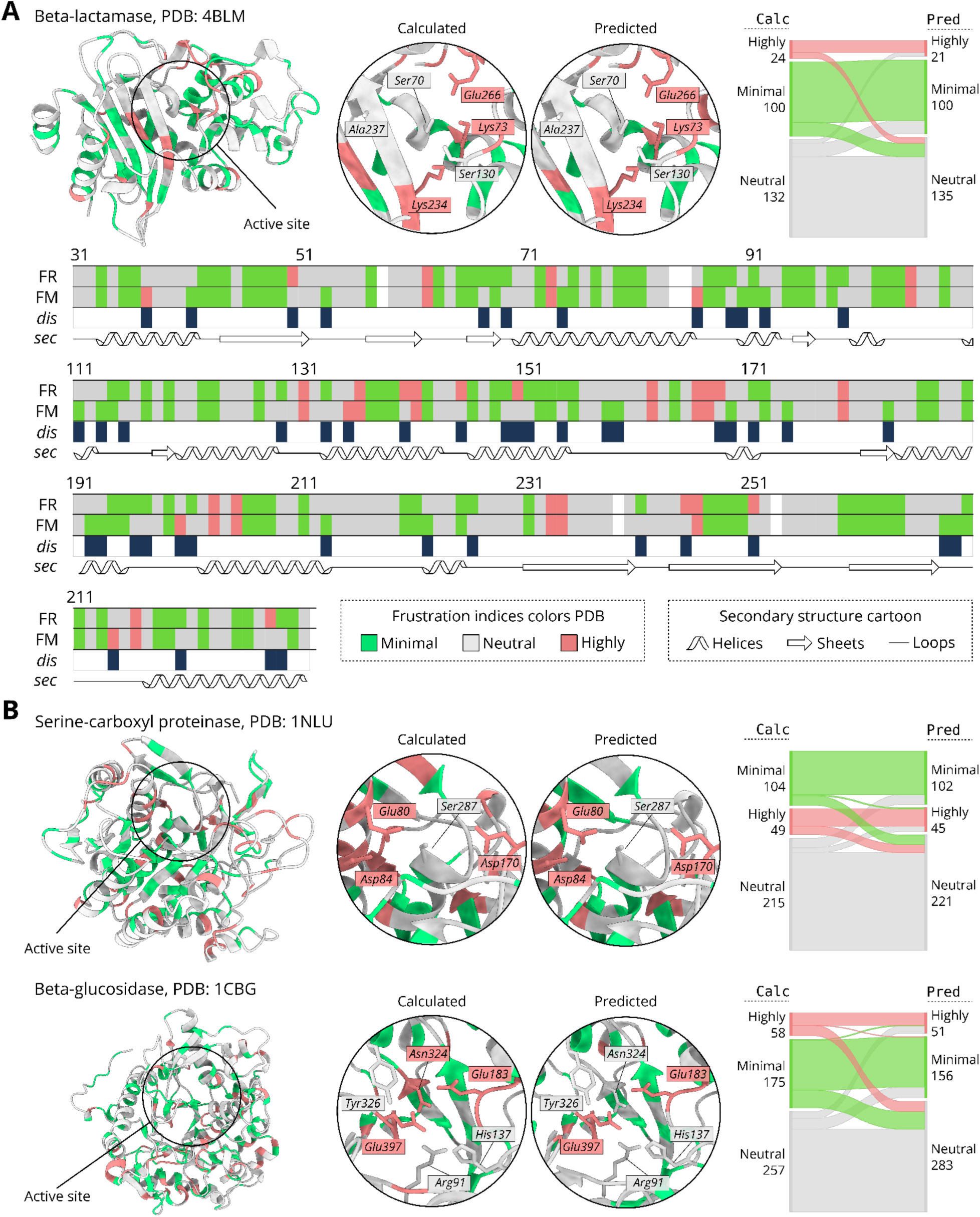
Test cases for the prediction of local protein single-residue frustration in enzyme active sites. (A) (left) Model structure colored by the calculated single-residue frustration indices from FrustratometeR [7,16,17] and indication of the active site residues based on the annotation from [6] for beta-lactamase (PDB: 4BLM). (middle) Key active-site residues (Ser70, Lys73, Ser130, Lys234, Asn237, Glu266) are shown in detail for the calculated and predicted single-residue frustration indices, with annotations colored by predicted frustration category (grey - neutral, red - highly). (right) Sankey diagram quantifies the flow of residue counts between calculated and predicted states, showing high agreement (e.g., 21 out of 24 highly frustrated residues were predicted correctly). (bottom) The sequence map aligns the frustration profiles from FrustratometeR (*FR*, calculated) and FrustraMPNN (*FM*, predicted), with disagreement (*dis*) indicated by dark blue bars. White blank rectangles indicate structure gaps. Fourth row represent secondary structure (*sec*) elements calculated by Dssp from PyRosetta [47]. (B) Additional enzymatic examples for serine-carboxyl proteinase (PDB: 1NLU, top) and beta-glucosidase (PDB: 1CBG, bottom). For both enzymes, the 3D structure, zoom-ins of the active site defined by [30], and Sankey diagrams are shown. In both cases, the active sites contain highly and neutrally frustrated residues that are accurately captured by the prediction method. Sequence map profiles for both are shown in the Supplementary Figure S5.

A similar pattern can also be seen in the two other examples (Figure 3B), serine carboxyl protease and beta-glucosidase. In most cases, the predicted frustration state is consistent with the FrustratometeR calculations (only Asn324 in beta-glucosidase was calculated as highly frustrated and predicted as neutral). The key residues for the active centers were obtained from the Mechanism and Catalytic Site Atlas [30]. In beta-glucosidase, the shift from highly frustrated to neutral is greater, and some neutrally frustrated residues were also incorrectly predicted as highly frustrated. There is no clear pattern discernible in the distribution of misclassified residues with regard to secondary structure and buriedness. For serine-carboxyl proteinase, more residues in loops are incorrectly predicted, while in the case of beta-glucosidase, more residues in helices are incorrectly predicted. Buriedness does not appear to have any influence in this regard (Supplementary Figure S5 and Table S1)

### FrustraMPNN is orders-of-magnitude faster than existing methods

The most immediate and transformative advantage of FrustraMPNN is its computational efficiency, which removes the primary barrier to widespread adoption of comprehensive single-residue frustration analysis. We systematically benchmarked prediction times for FrustraMPNN on a single GPU against the current standard tool, FrustratometeR [17] operating in single-residue mode, across proteins ranging from 50 to 1,000 residues (Figure 4A). For small proteins of around 100 residues, where FrustratometeR must perform ∼2,000 individual frustration calculations, FrustraMPNN provides around 1,200-fold speedup compared to single-CPU calculations. This advantage increases dramatically with protein size: for proteins of approximately 300 residues, which require 6,000 frustration evaluations and 19h using the FrustratometeR, FrustraMPNN completes the entire analysis in ∼30 seconds. Parallelizing the FrustratometeR across 21 CPUs for one CPU per mutation, the computation still takes 80 times longer. For larger proteins with more residues (above 10,000 individual calculations), the speedup exceeds 100-fold compared to 21 CPUs, with FrustraMPNN maintaining sub-minute computation times (Supplementary Table S2).

**Figure 4.**
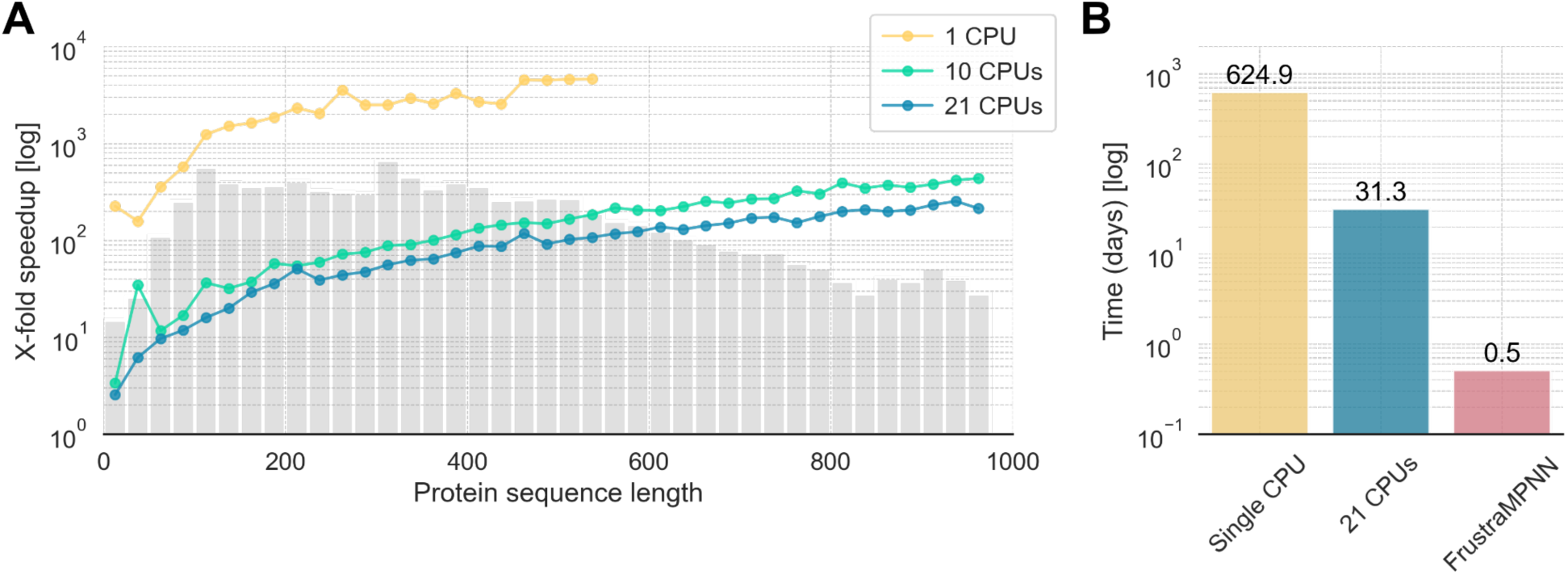
Computational acceleration achieved by FrustraMPNN for single-residue frustration analysis. (A) Mean speedup factor compared to FrustratometeR’s single-residue mode plotted against protein sequence length in 25-residue bins. Note that single-residue analysis requires 20 frustration calculations per position, one for each amino acid, making it ∼20× more computationally expensive than configurational or single-mutation analysis. Separate curves show FrustratometeR’s performance with 1 (yellow), 10 (green), and 21 (blue) CPU cores. Gray bars indicate the number of proteins in each size bin from the Alphafold validation set. FrustraMPNN achieves >100× speedup for proteins of ∼500 residues (10,000 individual calculations), with acceleration increasing almost linearly with protein size. (B) Estimated computational time for a complete single-residue frustration analysis of the *E. coli* proteome (∼4,300 proteins): 1.71 CPU-years with FrustratometeR in a single CPU vs. 12 GPU-hours with FrustraMPNN.

This scalability is particularly crucial for proteome-scale analyses. As an exemplar case, we analyzed the complete *E. coli* proteome (∼4,300 proteins) in single-residue mode, which would require approximately 1.71 CPU-years with the FrustratometeR, but can be completed in less than 12 GPU-hours with FrustraMPNN, thus corresponding to a 7,000-fold reduction in computational time (Figure 4B). The results demonstrate exceptional scalability advantages for our deep learning approach, particularly significant given that single-residue analysis requires evaluating all 20 amino acids at each position. Thus, FrustraMPNN predicts all single-residue frustration indices for all positions of whole proteomes in a single shot with great accuracy.

## Discussion

### Dataset diversity drives model generalizability

Three key factors emerged as critical for achieving robust generalization of FrustraMPNN across different protein topologies, secondary structure compositions and sizes: structural diversity in the training data, balanced representation of frustration categories, and incorporation of evolutionary information (Table 1 and Supplementary Table S3).

The importance of structural diversity is evident from the comparative analysis of the two training datasets. FireProt proteins exhibit a wide range of sizes (mean 189±95 residues), secondary structure compositions (31% helix, 28% sheet, 41% loop), and fold classes that represent major structural and evolutionary categories (SCOP, Structural Classification of Proteins [31]). In contrast, Megascale proteins are predominantly small (mean 56±12 residues), abundant in helical structures (52% helix, 18% sheet, 30% loop) and with limited fold diversity (Figure 1B and Supplementary Table S3). This difference in coverage of protein structural space directly correlates with generalization performance on the diverse AlphaFold human proteome validation set.

Balancing the categories of frustration during training proved essential for accurate prediction across the full range of frustration indices. Natural datasets exhibit a strong bias toward minimally frustrated residues (typically 40% of positions), with highly frustrated sites comprising only around 10% [2]. The calculated values for the FireProt and Megascale datasets follow this distribution to a reasonable extent for the wild-type residues (Supplementary Table S3). However, despite these observations having been made in previous works on the analysis of local energetic frustration in natural proteins [1], FrustraMPNN models trained on such imbalanced data show reduced sensitivity to highly frustrated regions, precisely the sites of most significant biological interest. Our balancing strategy, which equalizes representation across categories through targeted subsampling, improved prediction accuracy for highly frustrated residues by 23% without sacrificing overall performance (Table 2), surpassing the performance of the model trained on the saturated dataset that reproduces the natural distribution of frustration categories.

The superior generalization ability of the FireProt model could be due to its exposure to greater structural diversity during training. Although Megascale contains nearly 80 times more data points, its focus on small, artificially generated proteins means that it covers only a limited portion of the conformational and chemical space occupied by natural proteins. In contrast, the FireProt dataset contains less data but covers a broader spectrum of protein folds, secondary structure compositions, and functional classes. This finding that carefully curated diversity outweighs sheer data volume provides valuable insights for the future development of biological prediction models.

Our finding that dataset sequence length and structural diversity drive generalizability more than sheer data volume provides important methodological insights for the growing field of deep learning for proteins. The superior performance of models trained on the structurally diverse FireProt dataset, despite its smaller size compared to Megascale, challenges the common assumption that more data always yields better models, as it has been also ascertained in deep learning models for protein expression [32] and protein engineering [33]. This principle, that thoughtful curation targeting coverage of relevant chemical and structural space outweighs raw dataset size, should inform future efforts in biological machine learning. The result also validates our model development strategy, providing confidence that FrustraMPNN will perform robustly on the full diversity of natural proteins.

### Accelerating local frustration analysis for biological discovery

FrustraMPNN represents a transformative advance in computational structural biology, removing the primary barrier that has restricted frustration analysis to specialized applications. By achieving orders-of-magnitude acceleration while maintaining high accuracy, our tool makes this powerful analytical technique accessible to the broader scientific community. The implications extend across multiple domains of protein science, from fundamental mechanistic studies to applied biotechnology. The dramatic speedup enables entirely new research paradigms previously beyond reasonable computational reach. The frustration mapping across whole proteomes can now systematically identify functional sites in thousands of proteins, revealing patterns invisible to traditional sequence- or structure-based approaches.

In protein engineering, high-throughput screening of designed variants for altered frustration patterns provides a novel optimization criterion complementary to stability and binding affinity in protein-protein interactions. In fact, recent studies of Proteolysis-targeting chimeras (PROTACs) bound to target proteins of interest for degradation show higher interface frustration at the protein-protein interface, suggesting that calculations of interface frustration can guide rational, structure-based approaches to PROTAC design [34]. For drug discovery, frustration analysis of protein-protein interfaces can identify malleable regions susceptible to minor molecule disruption, guiding the design of allosteric modulators [35,36].

FrustraMPNN could also facilitate the optimization of specific frustration features, such as core stability or high functional frustration at interfaces, during the exploration of mutation landscapes. Single-residue frustration metrics for mutations could serve as terms in multi-criteria evaluation functions alongside standard potentials, such as the Rosetta energy function [37], or guide sampling strategies, such as Monte Carlo simulations, to select sequences that strike a balance between thermodynamic stability and the necessary functional dynamics.

### Limitations

A structural limitation of FrustraMPNN is its restriction to frustration indices for individual residues, which effectively projects a complex interaction network onto a single number. This granularity may fail to reveal specific frustrations in pairwise interaction patterns that are crucial for protein function. For example, catalytic residues often appear minimally frustrated at the level of individual residues because opposing energetic interactions cancel each other out, and highly frustrated specific contacts with supporting residues in the environment mask this intrinsic frustration [6]. Future developments could extend the architecture to predict frustration across contact maps, enabling the resolution of these pairwise energetic conflicts that are currently averaged out.

In addition, it is essential to note that the predictions from FrustraMPNN are not fully consistent with the calculations using the FrustratometeR [7,16,17]. What FrustraMPNN does offer is a balance between accuracy and an enormous prediction speed to enable screening many proteins, or, as shown, entire proteomes, in a relatively short time and calculating the whole single-residue frustration landscape for every position and every mutation. Of course, accuracy also comes into play here, and a lower accuracy is expected especially in unstructured regions such as loops or in transitions between sheets/helices and loops. These errors likely stem from the embeddings used by ProteinMPNN [22], leading to high uncertainty in the overall prediction, especially for low-quality backbones.

One indication of this could be the prediction of the identity of the catalytic residues in the enzyme test case beta-glucosidase (PDB: 1CBG, Figure 3B bottom). A key residue, Asn234, is incorrectly classified as neutral in the prediction and as highly frustrated in the calculation by FrustratometeR. This could be because Asn234 is located at the transition from a β-sheet to a loop, thus compromising the accuracy of the prediction. Although the other residues annotated as catalytic by the Mechanism and Catalytic Site Atlas [30] are also located in loops, the structural and sequence-specific context could be more accurate for the ProteinMPNN embeddings. Accordingly, an extension of the general network, or of the functionality of FrustraMPNN, could be to output uncertainties, such as the network’s confidence in a prediction and the upper and lower limits it estimates for the predicted single-residue frustration of a specific mutation. This could allow users to perform a more accurate, but time-consuming analysis with FrustratometeR, especially for highly uncertain residues, and to compare it with the prediction to validate the classification. However, determining inaccuracy requires a different approach to the network architecture adopted herein and is thus interesting for future projects.

Despite these limitations, the intersection of FrustraMPNN with other computational tools promises powerful synergies. Integration with biomolecular structure prediction methods such as AlphaFold2 [19] and AlphaFold3 [20], as well as Boltz2 [38], enables frustration analysis of proteins lacking experimental structures, expanding coverage to entire proteomes. Combination with design tools such as ProteinMPNN [22] or RFDiffusion3 [39] could incorporate frustration as an explicit design criterion, creating proteins with tailored dynamic properties. Such integrations position FrustraMPNN as a foundational component in the emerging ecosystem of deep learning-driven protein engineering and design tools.

### Implications for understanding protein function and evolution

Beyond its immediate utility as a computational tool, FrustraMPNN enables new perspectives on fundamental questions in protein biology. The ability to rapidly analyze frustration across large protein families reveals how evolution has shaped the distribution of frustrated sites to achieve diverse functions from conserved scaffolds. Systematic analysis of orthologs shows that frustration patterns at functional sites are often more conserved than sequence identity [4], suggesting that evolution operates on the energetic landscape as much as on specific residues.

FrustraMPNN could also provide insights into the design principles that distinguish natural from artificial proteins. Analysis of computationally designed proteins reveals they typically exhibit lower overall frustration than natural counterparts, reflecting optimization for stability over function [1]. This observation suggests that introducing controlled frustration could enhance the functional capabilities of designed proteins [40], moving beyond the current paradigm of maximizing thermodynamic stability toward engineering dynamic, responsive systems resembling natural counterparts.

## Materials and Methods

### Dataset curation and preparation

Two primary datasets were utilized in this study, each processed in different ways to generate comprehensive single-residue frustration profiles.

The FireProt dataset [26] comprises experimentally-characterized protein stability mutants from 100 natural proteins, primarily derived from thermophilic organisms and proteins of known industrial relevance. The curated dataset from ThermoMPNN [23] includes 3,404 experimental data points with measured ΔΔG values, concentrated in 15 extensively studied proteins that account for 65% of all measurements. The curated FireProt dataset was generated for ThermoMPNN and was obtained from https://doi.org/10.5281/zenodo.8169289 [23,41]. Due to processing errors affecting specific residues of 1A2P, 1AYE, 1HZ6, 1LZ1, 1RN1, 1RTP, 5DFR, and 5PTI, some positions were excluded from the dataset because the automatic processing pipeline was unable to match them to the calculated single-residue frustration values. The default FireProt dataset used in this study was thus reduced to 3,308 mutations.

The Megascale dataset [27] contains systematic deep mutational scanning data for 90 natural and 109 designed mini-proteins, each with complete single-site saturation mutagenesis. The curated dataset from ThermoMPNN [23] includes 272,712 experimental measurements. The Megascale dataset, which was curated based on the ThermoMPNN article, was obtained from https://doi.org/10.5281/zenodo.7992926 [42]. For the Megascale dataset, we excluded all rows with the mutation type “wild-type” (wt), because it does not allow us to match a single-residue frustration value for a specific position. Therefore, 1,481 data points were excluded, reducing the default Megascale dataset used in this study to 271,231 mutations.

To create comprehensive training data for single-residue frustration prediction, we generated two additional versions of the FireProt and Megascale datasets: (a) saturated datasets created by calculating frustration values for all 20 amino acids (including the native residue) at each position in single-residue mode, yielding 306,021 data points for the saturated FireProt dataset and 878,860 data points for the saturated Megascale; (b) a balanced dataset, achieved by subsampling to equalize representation across frustration categories, reducing the dataset of frustration indices to 144,013 data points in the case of FireProt and 433,135 in the case of Megascale (Supplementary Figure S2). For the saturation and balancing process, which is computationally intensive (requiring around 2,000 CPU-hours), two assumptions were made. First, that complete saturation of mutations should create a complete frustration landscape for the respective data sets and thus improve prediction accuracy. Second, that the balancing strategy would serve to distribute the frustration categories evenly, given the strong imbalance in the respective neutral, minimal, and highly frustrated residues.

### Frustration calculations

The single-residue frustration data for all structures in the different datasets were generated using a parallelized version of FrustratometeR [7,16,17], termed frustrapy, to calculate local energetic frustration from protein structures. This implementation features interactive visualizations, parallel processing, and performance optimizations [43]. The single-residue frustration was calculated using default parameters, namely no electrostatics and a sequence distance of 12, which represents the minimum number of amino acids separating two residues along the polypeptide chain.

To classify residues as minimal, highly, or neutrally frustrated, we use cutoffs of minimal frustration > 0.55 and high frustration < -1, as defined in previous works [4].

### AlphaFold DB validation set based on the human proteome

The external validation dataset consists of 3,415 human protein structures from the AlphaFold Protein Structure Database (AlphaFold DB) [24,25]. Proteins were randomly selected based on high confidence predictions (mean pLDDT > 70) and length between 50 and 1,000 residues. A complete list can be found on Zenodo (https://doi.org/10.5281/zenodo.17978321).

For each protein, we calculated complete single-residue frustration profiles using the frustrapy [43] wrapper of FrustratometeR [17] in single-residue mode, evaluating all 20 amino acids at each position, a computation that required approximately 8,000 CPU-hours but provides the ground truth for assessing FrustraMPNN’s ability to reproduce these comprehensive frustration landscapes in seconds rather than hours.

### *E. coli* dataset selection for proteome-wide screening test

In addition, we downloaded prediction files (version: UP000000625_83333_ECOLI_v4) for the *E. coli* model-organism proteome [44] from the AlphaFold DB [24,25] without further processing, as the proteome was used only to test the method’s speed across an entire organism, and we trusted the pre-filtering by the AlphaFold team. The resulting single-residue data were therefore not evaluated in detail, as with the other datasets. As mentioned above, single-residue frustration values were calculated for all positions and all possible mutations to generate a whole frustration landscape for all proteins using the frustrapy [43] wrapper of FrustratometeR [17].

### Model architecture and implementation

FrustraMPNN uses the same architecture as ThermoMPNN [23] to predict frustration indices. It integrates the pretrained ProteinMPNN [22] with a downstream module for predicting raw single-residue frustration values. The ProteinMPNN component, a Graph Neural Network (GNN) comprising 3 encoder and decoder layers, takes an input protein structure to process its wild-type sequence and the Gaussian radial basis functions (RBF)-encoded distances between backbone atoms (N, Cα, C, O) and a virtual Cβ for the target residue and its 48 nearest neighbors. Rather than generating sequence probabilities as in ProteinMPNN, FrustraMPNN extracts 128-dimensional decoder embeddings for the mutation site, which inherently capture local structural context via message passing during encoding. These structural embeddings are concatenated with the corresponding sequence embedding in the decoder layers to serve as input for the Frustration module. As in the ThermoMPNN architecture, this module applies a Light Attention block to refine the feature vector through self-attention, followed by a two-layer multilayer perceptron (MLP) that outputs the final frustration score. In contrast to ThermoMPNN, the predicted frustration value is not subtracted from the wild-type value to obtain single-residue frustration values, as in FrustratometeR [17].

For some experiments, ESM embeddings were incorporated as additional learning features using the pretrained ESM-2 t33 650M language model [29] available on GitHub (https://github.com/facebookresearch/esm). Similar to how ProteinMPNN structural embeddings were added, the ESM model processed the sequence for each structure. The resulting sequence embeddings were concatenated with the structural features, and the MLP was trained to predict continuous frustration indices.

### Training procedures

Different versions of the FrustraMPNN model were trained with the deep learning framework PyTorch Lightning [45] using the Adam optimizer with an initial learning rate of 0.001, reduced by a factor of 0.5 upon a validation loss plateau, using the ReduceLROnPlateau module from PyTorch [46]. Training proceeded for 100 epochs with a batch size of 32, employing gradient clipping with a norm of 1.0. Data augmentation included random rotation of input structures and dropout of 10% of edges during training. Data splitting was performed at the PDB level to prevent leakage, with 60% for training, 20% for validation, and 20% for testing.

For 5 random seeds, PDB IDs (with their associated mutation counts) were randomly ordered and then iteratively assigned to the training, validation, and subsequent test sets until their respective target mutation counts were approximated to guarantee that all mutations from a single PDB ID are allocated to only one subset. One seed represents one final dataset split. This PDB ID-level stratified splitting procedure was applied to both the FireProt and Megascale datasets, encompassing their respective default, saturated, and balanced versions.

Early stopping based on validation loss prevented overfitting, typically occurring around epoch 60-70. The training procedure was performed for all generated datasets from FireProt and Megascale (default, saturated, and balanced versions; see Methods section *Dataset curation and preparation*).

### Structural analysis

To identify residue location (surface, boundary, core) and secondary structural elements (helix, sheet, loop) in the different datasets, PyRosetta was used [47] with the LayerSelector (location) and the DsspMover (secondary structure).

### Computational benchmarking

Computational speed comparisons were performed on a dedicated compute node at Leipzig University with 2x AMD(R) EPYC(R) 7713 @ 2.0GHz - Turbo 3.7GHzX processors (64 cores each) and an NVIDIA A30V100 GPU. FrustraMPNN inference used a single GPU with batch size 1, while FrustratometeR calculations were run with 1, 10, or 21 CPU cores. Timing measurements excluded I/O operations and were averaged over 10 runs for each protein. Memory usage was monitored using system profilers to ensure fair comparison.

### Software and data availability

A CPU-parallelizable version of FrustratometeR [17] was implemented in-house using a Python wrapper, termed frustrapy [43], to facilitate the generation of local energetic single-residue frustration data for the different protein structure datasets. FrustraMPNN is implemented in Python 3.8+ with dependencies including PyTorch 1.12+, PyTorch Geometric 2.0+, and NumPy. The software is distributed under the MIT license on GitHub (https://github.com/schoederlab/frustraMPNN/) and includes comprehensive documentation, installation instructions, and tutorials. A Google Colab notebook provides immediate access without local installation. All training data, model weights, UniProt IDs, and validation results are deposited in Zenodo (https://doi.org/10.5281/zenodo.17978321).

## Supporting information

Supplementary informations

## Acknowledgment

We thank colleagues for helpful discussions and feedback on the manuscript. Leipzig University Computing Center provided computational resources. We acknowledge the developers of PyTorch, PyTorch Geometric, and the ESM-2 model for making their software freely available. We would like to thank the Google DeepMind team for sharing their protein structure predictions. Additionally, we would like to thank the ProteinMPNN and ThermoMPNN authors for making their tools freely available to the community.

## Author Contributions

Conceptualization: M.B., F.E., C.A.R-S. and J.M. Data Curation, M.B. and C.A.R-S. Formal analysis: M.B., F.E., R.G.P. and C.A.R-S. Funding acquisition: J.M., C.T.S., R.G.P. and C.A.R-S. Investigation: M.B., F.E. and C.A.R-S. Methodology: M.B., F.E., R.G.P. and C.A.R-S. Project administration: J.M., C.T.S. and C.A.R-S. Software: M.B. Supervision: C.A.R-S. and J.M. Validation: M.B. and F.E. Visualization: M.B. and F.E. Writing—original draft: M.B., F.E. and C.A.R.-S. Writing—review and editing: M.B., F.E., R.G.P., C.T.S., C.A.R.-S. and J.M.

