## Supplementary informations for "FrustraMPNN: An ultra-fast deep learning tool for proteome-scale analysis of deep mutational single-residue local energetic frustration in proteins"

### Supplementary Figures

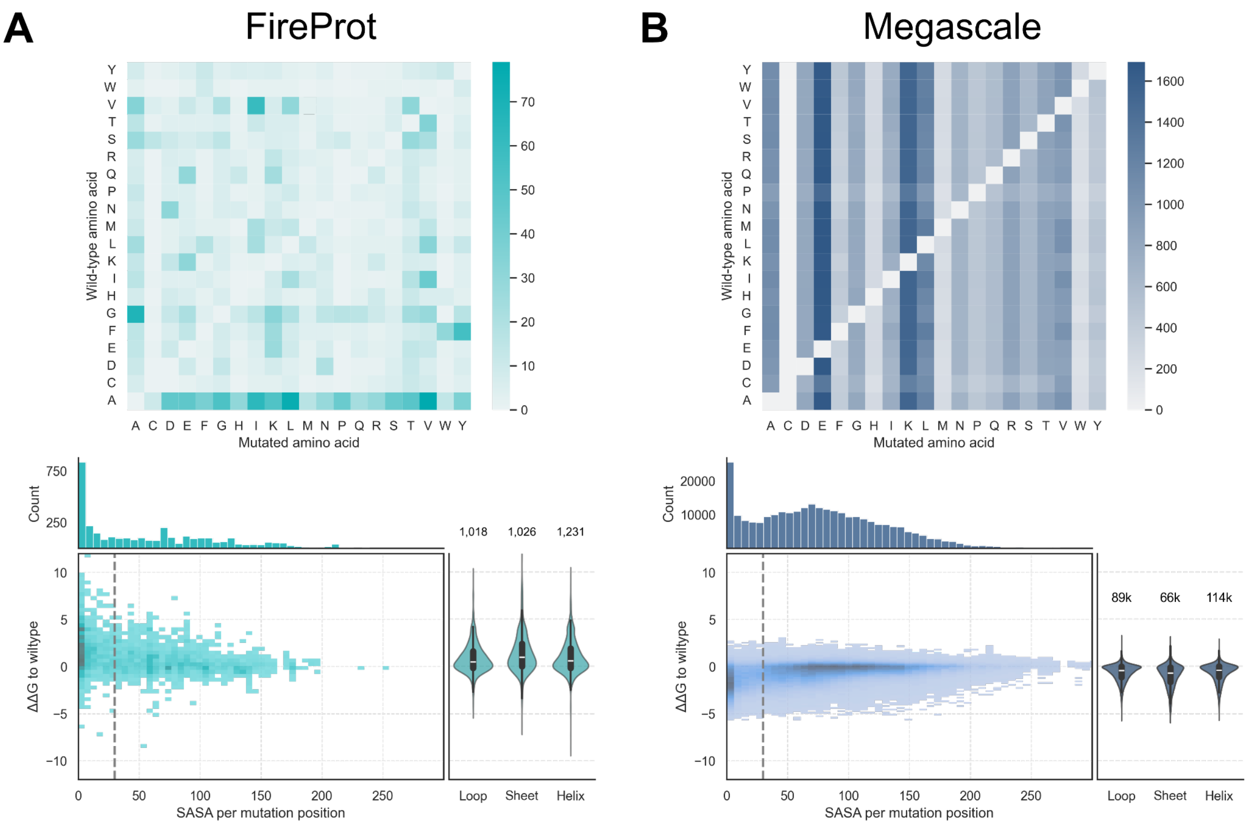


**Figure S1. Detailed dataset comparison.** Amino acid substitution matrices showing mutation patterns in each dataset (top) and distribution of mutation-induced stability changes (ΔΔG) against Solvent Accessible Surface Area (SASA) (bottom) for the default (a) FireProt and (b) Megascale datasets. Each bottom panel includes: (Top) SASA distribution of mutated residues; (Center) 2D histogram of ΔΔG versus SASA; (Right) violin plots of ΔΔG distributions conditioned on secondary structure (loop, sheet, helix; total counts indicated above, k = thousand). Dashed vertical lines represent the SASA cutoff value of 30 for the distinction between core/boundary and surface.

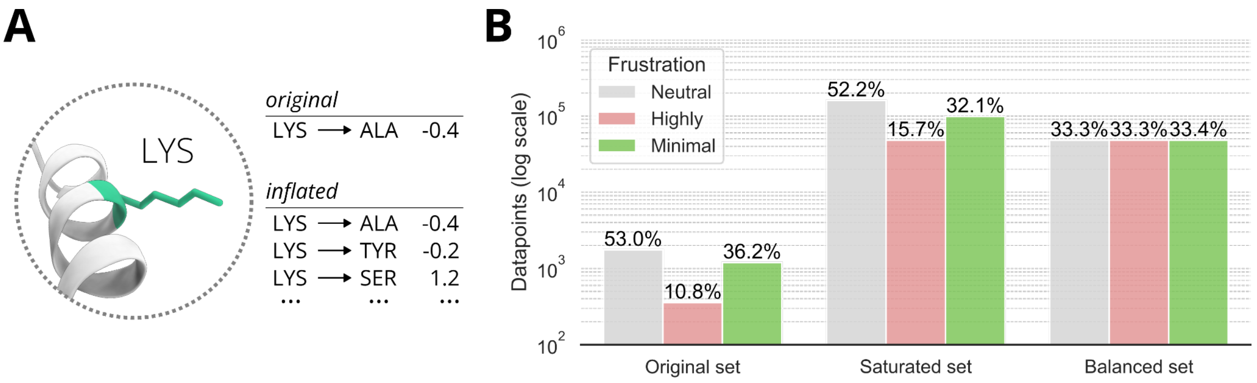


**Figure S2. FireProt dataset inflation as an example of data augmentation**. (A) Data points for the saturation example: calculate single-residue frustration values for every possible other amino acid at a specified position. This procedure was performed for all positions across all structures in the datasets. (B) Bar plot shows the distribution of data points across the frustration categories for the default, saturated, and balanced sets as an example of the FireProt dataset.

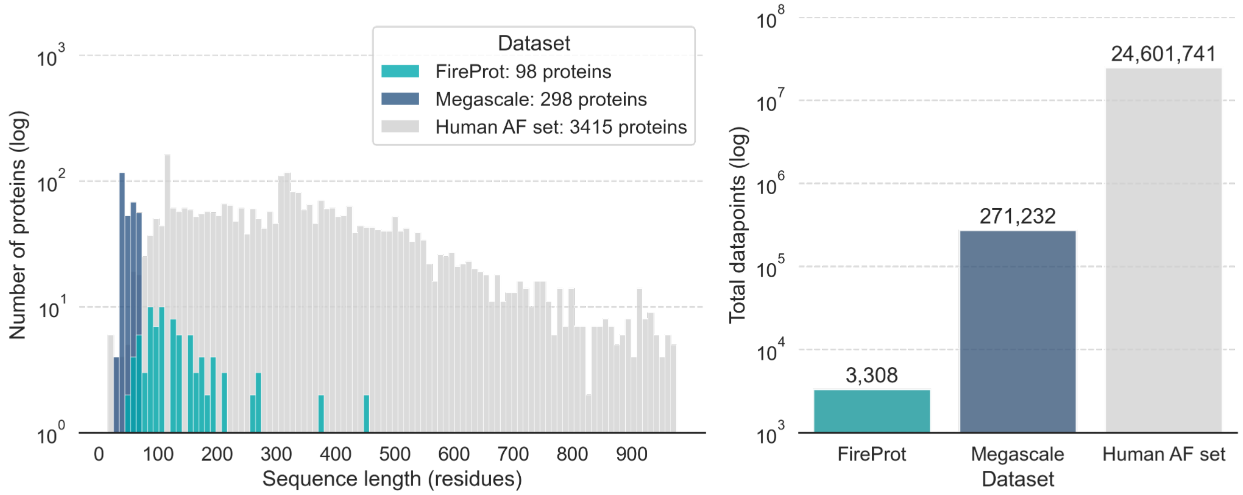


**Figure S3. Comparison of the FireProt, Megascale, and the external AlphaFold validation set based on the human proteome.** The AlphaFold validation set was obtained from the AlphaFold DB and filtered to structures with mean pLDDT > 70. The number of data points (right) is based on the default FireProt and Megascale sets, and for AlphaFold, it is the total number of residues for all proteins multiplied by 20 for the single-residue frustration calculation.

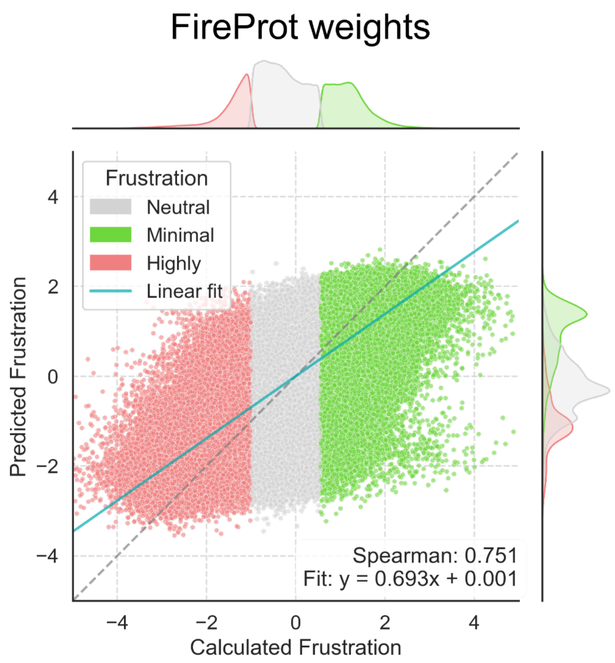


**Figure S4. Prediction scatterplot for randomly selected proteins.** Correlation between calculated and predicted frustration indices for 100 randomly selected validation proteins from the AlphaFold human proteome set, demonstrating consistent performance across the full range of frustration values. Points are colored based on the calculated frustration indices. Cutoffs for categories are > 0.55 for minimal frustration and < -1 for high frustration.

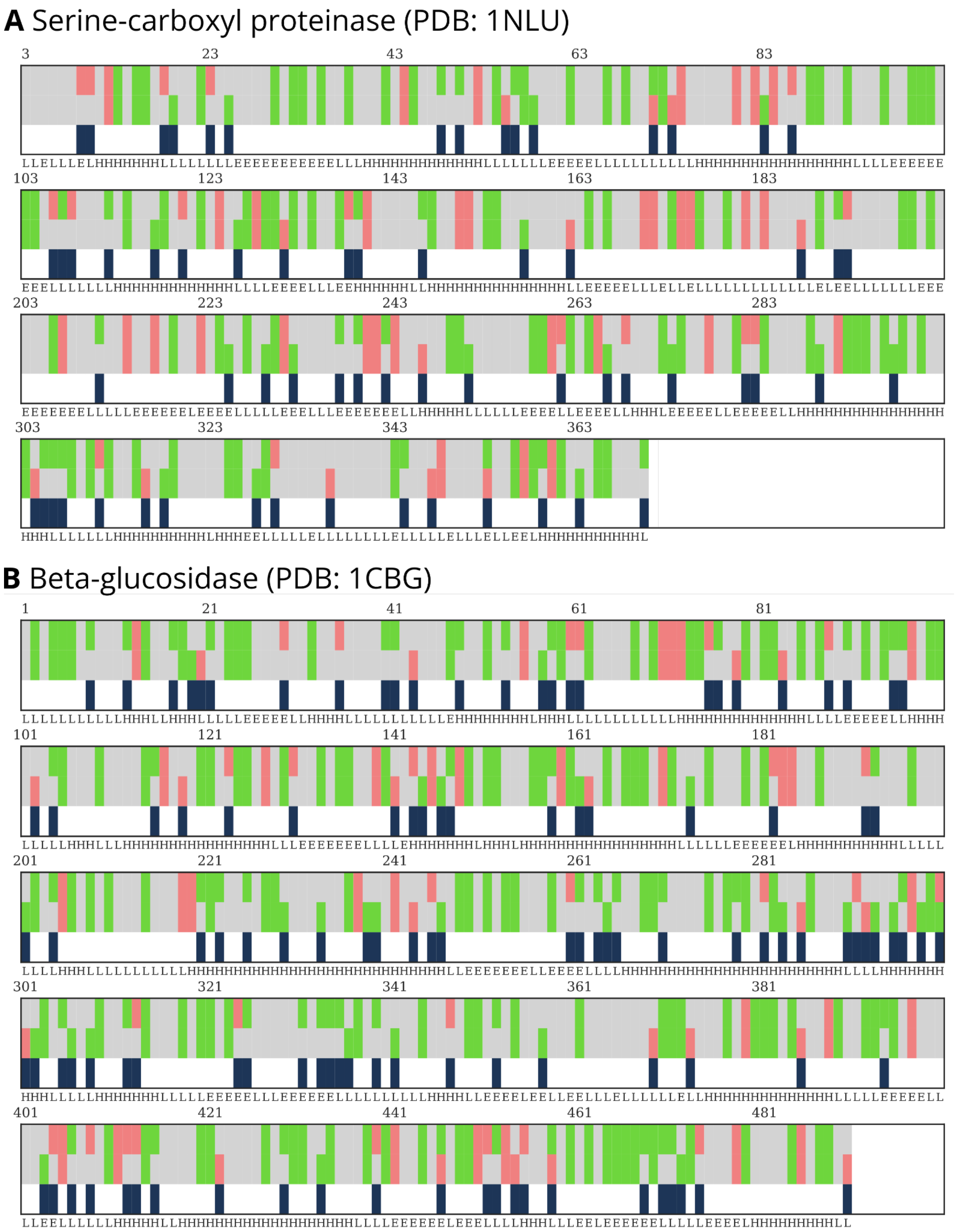


**Figure S5. Sequence maps for the chosen examples 1NLU and 1CBG.** The first row of every block always corresponds to the calculated single-residue frustration values, the second to the predicted ones. The third row indicates disagreement between the calculation and prediction. The fourth row represents the secondary structure elements (H - Helix, E - Sheet, L - Loops) calculated by Dssp from PyRosetta [1].

### Supplementary Tables

**Table S1. Prediction errors for the three chosen test cases broken down by secondary structure and buriedness.** Raw count of misclassified residues for the chosen test enzymes beta-lactamase (PDB: 4BLM), serine-carboxyl proteinase (PDB: 1NLU), and beta-glucosidase (PDB: 1CBG). Misclassified residues are further split into their location in secondary structure elements and their buriedness (value in parentheses). A solvent accessible surface area (SASA) value above 30 Å$^2$ was chosen for surface residues.

| | | **4BLM** | **1NLU** | **1CBG** |
|---|---|---|---|---|
| Length | | 261 | 369 | 491 |
| Incorrect classified (SASA > 30) | Total | 49 (27) | 64 (30) | 119 (59) |
| | Helix | 24 (12) | 17 (8) | 56 (27) |
| | Sheet | 5 (0) | 16 (5) | 17 (8) |
| | Loop | 20 (15) | 31 (17) | 46 (24) |

**Table S2.** Mean inference time and single-residue calculation time for the external AlphaFold validation set, a random selection from the human proteome between 25 and 1,000 amino acids in length (see Materials and Methods). The proteins were binned into 25-residue bins. All times are measured in milliseconds and represent mean values for the bin.

| **CPU count** | **Bin** | **Inference time (ms)** | **Single residue calculation** | **Speedup** |
|---|---|---|---|---|
| 1 CPU | 37.5 | 21434 | 3388948 | 158.11 |
| 1 CPU | 62.5 | 11857 | 4255774 | 356.26 |
| 1 CPU | 87.5 | 20363.11 | 7941105.22 | 575.34 |
| 1 CPU | 112.5 | 14287.75 | 17197309.74 | 1244.66 |
| 1 CPU | 137.5 | 17676.44 | 23726239.78 | 1517.19 |
| 1 CPU | 162.5 | 20711.73 | 26728891 | 1633.72 |
| 1 CPU | 187.5 | 19004.17 | 34156270.17 | 1871.02 |
| 1 CPU | 212.5 | 24513.06 | 45543815.72 | 2327.55 |
| 1 CPU | 237.5 | 23417.8 | 42730463.9 | 2052.7 |
| 1 CPU | 262.5 | 19398.75 | 69083346.5 | 3567.26 |
| 1 CPU | 287.5 | 28328.25 | 66320242.5 | 2509.17 |
| 1 CPU | 312.5 | 38508.3 | 68407985.1 | 2508.83 |
| 1 CPU | 337.5 | 26938.7 | 71870640.3 | 2936.57 |
| 1 CPU | 362.5 | 42564.29 | 81013808.43 | 2592.73 |
| 1 CPU | 387.5 | 27583.63 | 86730141.75 | 3325.69 |
| 1 CPU | 412.5 | 34936.17 | 88403998.5 | 2692.55 |
| 1 CPU | 437.5 | 35803 | 92219462 | 2575.75 |
| 1 CPU | 462.5 | 27473 | 124850049.3 | 4550.74 |
| 1 CPU | 487.5 | 28130.25 | 126932077.8 | 4520.25 |
| 1 CPU | 512.5 | 30161.2 | 139574076.8 | 4618.91 |
| 1 CPU | 537.5 | 31803 | 147532612 | 4642.88 |
| 10 CPUs | 12.5 | 11870.33 | 39375.83 | 3.37 |
| 10 CPUs | 37.5 | 16941.25 | 1452421.38 | 34.6 |
| 10 CPUs | 62.5 | 14449.51 | 194186.02 | 11.72 |
| 10 CPUs | 87.5 | 19235.41 | 340835.51 | 16.95 |
| 10 CPUs | 112.5 | 124129.66 | 1016486.29 | 36.4 |
| 10 CPUs | 137.5 | 141608.63 | 659586.64 | 31.78 |
| 10 CPUs | 162.5 | 21083.26 | 813562.59 | 37.52 |
| 10 CPUs | 187.5 | 83828.1 | 1604624.8 | 57.62 |
| 10 CPUs | 212.5 | 86322.57 | 1400702.43 | 54.42 |
| 10 CPUs | 237.5 | 22272.76 | 1379257.72 | 59.57 |
| 10 CPUs | 262.5 | 23968.17 | 1863139.29 | 72.2 |
| 10 CPUs | 287.5 | 23421 | 1798748.15 | 75.4 |
| 10 CPUs | 312.5 | 26542.09 | 2524588.78 | 88.57 |
| 10 CPUs | 337.5 | 25761.88 | 2270492.66 | 90.58 |
| 10 CPUs | 362.5 | 26562.91 | 2591237.37 | 100.64 |
| 10 CPUs | 387.5 | 26561.68 | 3046465.28 | 115.01 |
| 10 CPUs | 412.5 | 28000.25 | 3950869.88 | 134.05 |

| CPU count | Bin | Inference time (ms) | Single residue calculation | Speedup |
|---|---|---|---|---|
| 10 CPUs | 437.5 | 31054.7 | 4365821.28 | 145.61 |
| 10 CPUs | 462.5 | 28168.29 | 4365889.51 | 153.12 |
| 10 CPUs | 487.5 | 32261.77 | 4684409.26 | 148.81 |
| 10 CPUs | 512.5 | 32661.25 | 5259986.96 | 167.02 |
| 10 CPUs | 537.5 | 34709.87 | 6177031.07 | 185.03 |
| 10 CPUs | 562.5 | 38688.95 | 7898994.43 | 217.44 |
| 10 CPUs | 587.5 | 38587.54 | 7512155.31 | 205.82 |
| 10 CPUs | 612.5 | 39469.39 | 7726015.5 | 203.92 |
| 10 CPUs | 637.5 | 40567.42 | 8707847.61 | 223.74 |
| 10 CPUs | 662.5 | 39342.1 | 9922767.08 | 253.54 |
| 10 CPUs | 687.5 | 40491.23 | 9765664.67 | 243.31 |
| 10 CPUs | 712.5 | 43355.36 | 11652294.6 | 267.65 |
| 10 CPUs | 737.5 | 40791.19 | 10989793.46 | 270.28 |
| 10 CPUs | 762.5 | 41624.39 | 13697252.33 | 326.42 |
| 10 CPUs | 787.5 | 47444.89 | 14007465.16 | 303.49 |
| 10 CPUs | 812.5 | 49028.21 | 18652272.43 | 393.7 |
| 10 CPUs | 837.5 | 50915.64 | 17107988.09 | 347.43 |
| 10 CPUs | 862.5 | 49647.47 | 18031249.87 | 372.9 |
| 10 CPUs | 887.5 | 50571.36 | 17861533.71 | 353.41 |
| 10 CPUs | 912.5 | 56415 | 21238899.9 | 381.28 |
| 10 CPUs | 937.5 | 59741.43 | 23958587.93 | 419.48 |
| 10 CPUs | 962.5 | 52152.73 | 22659186.36 | 438.88 |
| 21 CPUs | 12.5 | 10353.67 | 26513.17 | 2.56 |
| 21 CPUs | 37.5 | 15778.08 | 94254.25 | 6.2 |
| 21 CPUs | 62.5 | 16906 | 157487.2 | 9.71 |
| 21 CPUs | 87.5 | 16050.97 | 187859.89 | 11.85 |
| 21 CPUs | 112.5 | 16343.06 | 260637.64 | 16 |
| 21 CPUs | 137.5 | 18365.9 | 371501.26 | 19.98 |
| 21 CPUs | 162.5 | 18598.01 | 632976.77 | 29.21 |
| 21 CPUs | 187.5 | 20360.95 | 752843.74 | 35.82 |
| 21 CPUs | 212.5 | 20988.2 | 1175103.35 | 50.96 |
| 21 CPUs | 237.5 | 20954.61 | 805547.81 | 39 |
| 21 CPUs | 262.5 | 23614.26 | 1096559.79 | 44.11 |
| 21 CPUs | 287.5 | 23515.07 | 1153502.22 | 47.52 |
| 21 CPUs | 312.5 | 26110.2 | 1446611.99 | 56.2 |
| 21 CPUs | 337.5 | 24486.18 | 1562528.97 | 62.5 |
| 21 CPUs | 362.5 | 29542.32 | 1898491.29 | 64.74 |
| 21 CPUs | 387.5 | 29927.68 | 2205683.48 | 74.68 |
| 21 CPUs | 412.5 | 28529.78 | 2519831.3 | 87.5 |
| 21 CPUs | 437.5 | 29457.09 | 2517553.78 | 86.89 |
| 21 CPUs | 462.5 | 31229.65 | 3488128.34 | 117.73 |
| 21 CPUs | 487.5 | 32322.83 | 2975903.04 | 92.07 |
| 21 CPUs | 512.5 | 31324.49 | 3192922.57 | 102.5 |
| 21 CPUs | 537.5 | 35778.39 | 3776868.49 | 107.82 |

| CPU count | Bin | Inference time (ms) | Single residue calculation | Speedup |
|---|---|---|---|---|
| 21 CPUs | 562.5 | 35595.41 | 4232197.78 | 117.65 |
| 21 CPUs | 587.5 | 36965.31 | 4611158.4 | 123.02 |
| 21 CPUs | 612.5 | 38833.68 | 5307342.84 | 137.95 |
| 21 CPUs | 637.5 | 39525.33 | 5016036.21 | 130.08 |
| 21 CPUs | 662.5 | 39154.95 | 5571746.88 | 142.46 |
| 21 CPUs | 687.5 | 38872.76 | 5853028.42 | 150.56 |
| 21 CPUs | 712.5 | 40554 | 6881685.58 | 170.4 |
| 21 CPUs | 737.5 | 44266.83 | 7556704.97 | 174.32 |
| 21 CPUs | 762.5 | 51002.44 | 7562586.33 | 152.18 |
| 21 CPUs | 787.5 | 44601.35 | 7898661.73 | 176.96 |
| 21 CPUs | 812.5 | 49889.18 | 9912563.65 | 199.67 |
| 21 CPUs | 837.5 | 46213.27 | 9654070.4 | 208.17 |
| 21 CPUs | 862.5 | 56361.05 | 11144131.42 | 199.35 |
| 21 CPUs | 887.5 | 54103.88 | 11022751.19 | 206 |
| 21 CPUs | 912.5 | 52034.83 | 12052038.33 | 233.11 |
| 21 CPUs | 937.5 | 53898.78 | 13771875.67 | 254.3 |
| 21 CPUs | 962.5 | 53239.38 | 11383228.31 | 214.05 |

83

**Table S3. Dataset characteristics for FireProt and Megascale influencing model generalizability**. Comparison of structural and sequence diversity between training datasets and fraction for minimal, neutral, and highly frustrated residues (cutoff minimal > 0.55 frustration index [FI], highly < -1.0 FI). For the first frustration categories (marked with *), only the frustration of individual amino acids of the wild-type sequence was counted; for the second (marked with +), the frustration of the mutations of individual residues was counted.

| Dataset Property | FireProt | | Megascale | |
|---|---|---|---|---|
| Number of proteins | 98 | | 298 | |
| Protein size (mean ± std) | 189 ± 95 | | 56 ± 12 | |
| Size range (residues) | 45-495 | | 35-75 | |
| **Secondary Structure Composition** | | | | |
| α-helix (%) | 31 | | 52 | |
| β-sheet (%) | 28 | | 18 | |
| Loop/coil (%) | 41 | | 30 | |
| **Mutation Distribution** | | | | |
| Mutations per protein (median) | 18 | | 1,330 | |
| Coverage of amino acids (%) | 95 | | 100 | |
| Alanine mutations (%) | 24.4 | | 5.3 | |
| **Frustration categories wildtype (*)** | | | | |
| Minimal (%) | 36.8 | | 39.4 | |
| Neutral (%) | 52.3 | | 49.5 | |
| Highly frustrated (%) | 10.9 | | 11.1 | |
| **Frustration categories datasets (+)** | | | | |
| | *Default set* | *Saturated set* | *Default set* | *Saturated set* |
| Minimal (%) | 36.2 | 32.1 | 29.7 | 28.9 |
| Neutral (%) | 53.0 | 52.2 | 55.0 | 54.6 |
| Highly frustrated (%) | 10.8 | 15.7 | 15.2 | 16.4 |
